# UniFlow: Unifying protein conformational ensemble generation and machine-learned force fields with a scalable normalizing Flow

**DOI:** 10.64898/2026.07.17.739266

**Authors:** Yikai Liu, Ming Chen, Guang Lin

## Abstract

Molecular dynamics (MD) provides a principled method for modeling equilibrium protein conformational energy landscapes, but its computational cost limits access to long timescales and larger protein systems. Recently, generative protein ensemble models and machine-learned coarse-grained force fields have emerged as complementary approaches for accelerating conformational sampling. However, they are typically developed separately despite modeling the same underlying equilibrium distribution. We introduce **UniFlow**, the first scalable generative model that unifies protein ensemble generation and machine-learned coarse-grained force fields for molecular dynamics simulation within a single framework. UniFlow employs an internal-coordinate normalizing flow that supports efficient i.i.d. sampling, exact likelihood evaluation, and differentiable energy and force computation. Across diverse protein systems, UniFlow generates ensembles that closely match reference MD simulations, generalizes to proteins beyond its training dataset, and samples substantially faster than diffusion-based ensemble-generation baselines. The same learned density further enables stable long-timescale molecular dynamics simulations. Together, UniFlow paves the way for a unified class of models that bridges generative ensemble modeling with physics-based molecular simulation.

**Code:** https://github.com/Harrydirk41/UniFlow.git

## 1 Introduction

Proteins function through ensembles of interconverting conformations rather than a single static structure. Their conformational distributions govern major biomolecular processes, including molecular recognition, catalysis, and allostery Whisstock & Lesk (2003). All-atom molecular dynamics (MD) simulations provide a principled route to estimating such equilibrium ensembles Shaw et al. (2010); Robustelli et al. (2018), but their ability to capture rare transitions and long-timescale behavior is often limited by computational cost.

Recent advances in machine learning have opened two major directions for addressing this challenge. The first is generative protein ensemble modeling, which aims to learn transferable protein conformational energy landscapes directly from large-scale structural and simulation data, typically through likelihood-based objectives Lu et al. (2023); Lewis et al. (2025); Jing et al. (2024a); Noé et al. (2019); Zheng et al. (2024); Wayment-Steele et al. (2024); Jing et al. (2024b); Raja et al. (2025); Lelièvre et al. (2023); Du et al. (2024); Fu et al. (2022); Arts et al. (2023); Liu et al. (2025; 2026). The second is machine-learned coarse-grained force fields (ML-CGFFs) Durumeric et al. (2026); Charron et al. (2025); Majewski et al. (2023), which aim to construct physically consistent effective potentials on coarse-grained representations. These directions are closely related thermodynamically, as both seek to represent equilibrium distributions over protein conformations. However, they have largely been developed independently: generative models emphasize scalable distribution learning and efficient sampling, whereas ML-CGFFs emphasize physical consistency and accurate force predictions for long-scale dynamical simulations.

Motivated by the connection and the current gap between the two research directions, we introduce **UniFlow**, a scalable normalizing-flow framework that unifies protein backbone ensemble generation with ML coarse-grained force-field learning. To our knowledge, UniFlow is the first scalable and transferable architecture designed to support both objectives within a single model. UniFlow operates in protein internal-coordinate space and uses a coupling-based normalizing flow with circular spline transforms parameterized by geometry-aware Transformer conditioners. It is trained on large-scale equilibrium MD trajectories, enabling transferable modeling of protein conformational distributions. Unlike diffusion-based approaches, UniFlow provides exact likelihoods over protein conformations. This makes the learned distribution directly usable as a differentiable potential for molecular dynamics simulations.

Our contributions are:

1. We introduce **UniFlow**, the first scalable generative framework that unifies protein conformational ensemble generation with machine-learned force-field modeling. Built on a normalizing flow over protein internal coordinates, UniFlow supports efficient i.i.d. sampling, exact likelihood evaluation, and differentiable energy computation for molecular dynamics within a single model.
2. We develop a scalable training strategy for transferable generative modeling of protein conformational ensembles in internal-coordinate space. By combining likelihood-based learning with geometry-aware supervision, UniFlow mitigates the strong locality of internal coordinates and generalizes across protein systems of different sizes, including unseen sequences.
3. We propose a distillation recipe that transfers a pretrained UniFlow model into a lightweight, system-specific potential for efficient molecular dynamics. The distilled model enables stable Langevin simulations while substantially reducing the computational cost of force evaluation.

## 2 Background

### 2.1 Normalizing Flows

Normalizing flows are likelihood-based generative models that define an invertible and differentiable transformation between a target variable and a simple base distribution. For data *x* ∈ ℝ^*D*^, a flow specifies

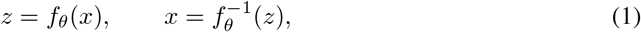

where *z* follows a tractable base density *p*_0_ such as a Gaussian distribution. The induced density is obtained through the change-of-variables formula,

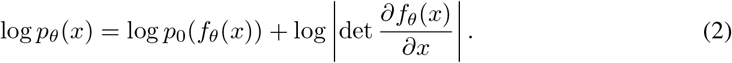

Maximum-likelihood training therefore minimizes

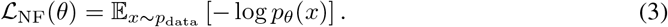

After training, exact samples are generated in one step by drawing *z* ~ *p*_0_ and evaluating 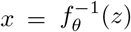.

The main architectural challenge is to construct expressive transformations while retaining tractable Jacobian determinants and efficient inverse maps. Coupling-based flows, such as RealNVP Dinh et al. (2016), achieve this by updating subsets of variables conditioned on the remaining variables. Autoregressive flows Papamakarios et al. (2017); Zhai et al. (2024) increase expressivity by allowing each variable or block to depend on an ordered set of preceding variables.

For angular internal coordinates, the target domain is not Euclidean. Because −*π* and *π* represent the same point, each angular variable lies on the one-dimensional torus T ≅ (−*π, π*], and a collection of *D* angles lies on T^*D*^. A periodic flow therefore uses an invertible map

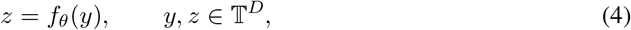

together with a periodic base density *p*_0_. To respect the periodic identification, the transformation must be continuous, bijective, and differentiable across the boundary. Neural spline flows Durkan et al. (2019) proposed circular rational-quadratic spline transformations, which provide expressive periodic maps while preserving exact likelihood evaluation.

### 2.2 Protein Internal-Coordinate Representation

A protein backbone can be parameterized by internal coordinates consisting of bond lengths *r*, valence angles 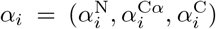, and torsion angles *τ*_*i*_ = (*ϕ*_*i*_, *ψ*_*i*_, *ω*_*i*_) for each residue *i* = 1, …, *L*. Because backbone bond lengths fluctuate over a substantially narrower range than the angular degrees of freedom, UniFlow learns only the joint distribution of the 6*L* angular coordinates (*τ, α*). For direct ensemble generation and geometry conditioning within the flow, bond lengths are set to their ideal values. When constructing the molecular-dynamics potential, the learned angular density is augmented with a stiff harmonic bond-length distribution, as described in Sec. 4.4. The internal-coordinate representation removes the global translational and rotational degrees of freedom and therefore does not require an *SE*(3)-equivariant invertible architecture. However, a local change in one internal coordinate can induce global Cartesian displacements across many atoms, making long-range structural dependencies difficult to capture.

To address this issue, we consider the native structure of a protein to provide informative local and global geometric context for modeling conformational ensembles. Native protein structures are widely available Jumper et al. (2021). We therefore use the native backbone torsions as a protein-specific reference for the more flexible torsional degrees of freedom. In contrast, backbone valence angles fluctuate within relatively narrow ranges around well-defined equilibrium values, so we use ideal valence angles as a common reference. Accordingly, rather than modeling absolute angles directly, we represent each conformation using angular displacements

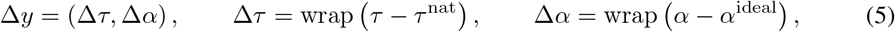

where *τ* ^nat^ denotes the backbone torsions of the native structure, *α*^ideal^ denotes the equilibrium valence angles, and wrap(·) maps angular differences to (−*π, π*].

## 3 Method

UniFlow models the conditional distribution

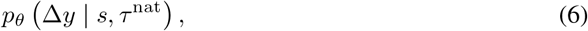

where *s* denotes the amino-acid sequence, *τ* ^nat^ denotes the native backbone torsions, and Δ*y* is the internal-coordinate displacement representation defined in Eq. 5.

A central challenge in internal-coordinate generative modeling is recovering global structural organization from locally defined angular variables. UniFlow addresses this challenge using a sequence- and native-conditioned normalizing flow composed of circular coupling layers. Within each layer, masked internal-coordinate displacements are reconstructed into a Cartesian backbone and encoded by an SE(3)-invariant geometric Transformer. A two-track Transformer then combines the encoded structural representation with the amino-acid sequence to predict the parameters of the circular coupling transformation. This construction preserves exact likelihood evaluation and efficient parallel generation while allowing each transformation to depend on the global geometry implied by the current layer internal coordinates. In addition to maximum-likelihood training, we apply auxiliary geometric objectives to generated structures to directly regularize global physical observables that are not explicitly targeted by the internal-coordinate likelihood. An overview of UniFlow is shown in Fig. 1.

**Figure 1.**
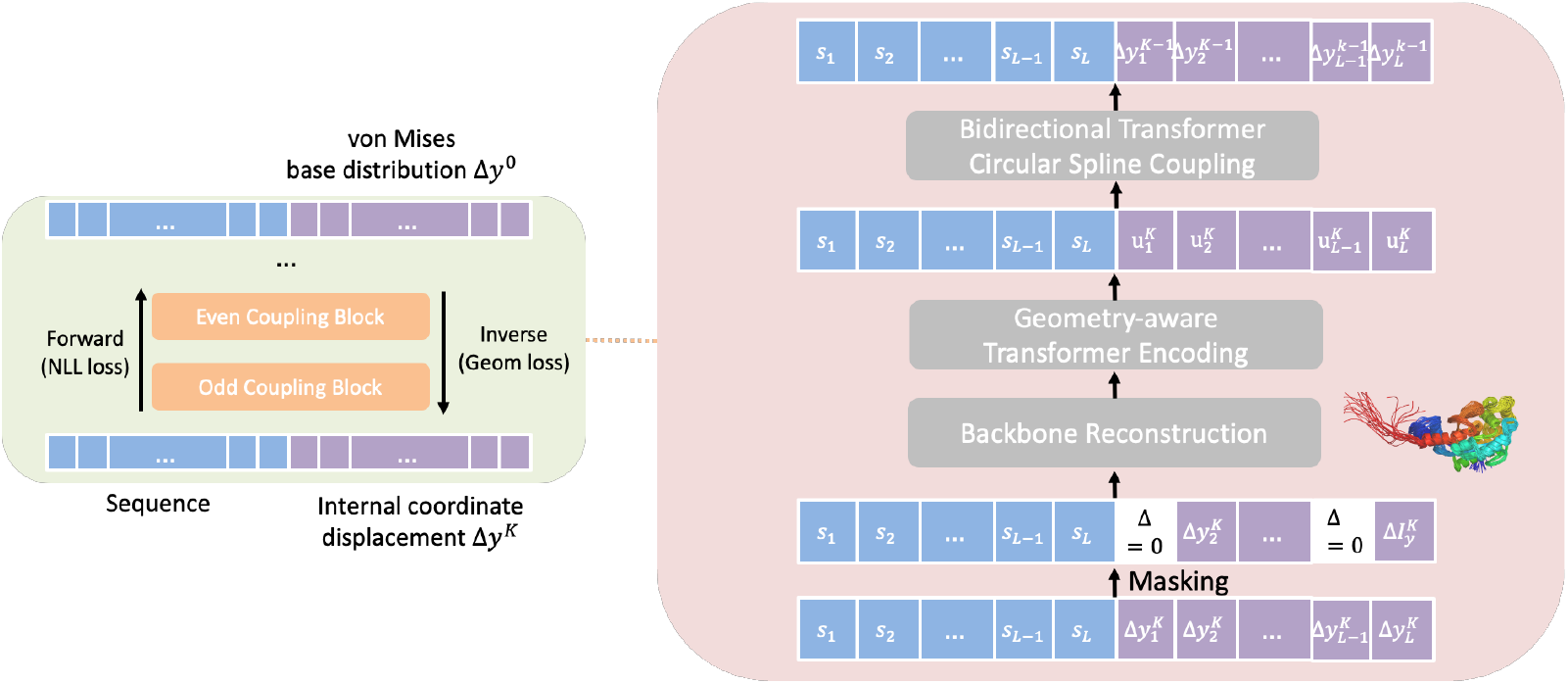
Overview of **UniFlow.** A protein backbone is represented using fixed ideal bond lengths and six angular degrees of freedom per residue: three backbone torsions and three valence angles. UniFlow models angular displacements from native torsions and ideal valence angles with a stack of invertible circular coupling layers. Each layer preserves one subset of residues under an odd– even masking scheme and transforms the complementary subset using periodic rational-quadratic splines. The spline parameters are predicted by a geometry-aware conditioner comprising three components: backbone reconstruction, a geometry-aware Transformer that encodes structure representations, and a two-stack sequence–structure Transformer. During generation, samples from a factorized von Mises base distribution are transformed through the flow into angular displacements and reconstructed into Cartesian backbone coordinates using NeRF.

### 3.1 Circular Coupling Flow

UniFlow uses a stack of *N* = 12 circular coupling layers. We denote the base-space variable by Δ*y*^(0)^ and the data-space variable by Δ*y*^(*N*)^ = Δ*y*. The generative transformation is

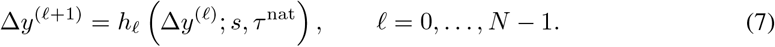

Thus, generation maps Δ*y*^(0)^ from the base distribution to the data-space displacement Δ*y*^(*N*)^, whereas likelihood evaluation applies the same layers in reverse.

The base density factorizes over valid residue-channel pairs:

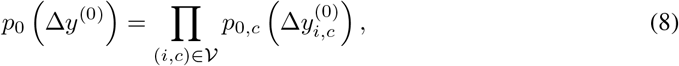

where *V* denotes the set of valid angular coordinates. Each channel follows a von Mises distribution,

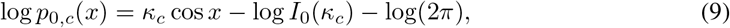

where *κ*_*c*_ is the concentration parameter for channel *c* and *I*_0_(·) is the modified Bessel function of the first kind.

At layer *ℓ*, we use an alternating odd–even residue mask. The active and inactive residue sets are

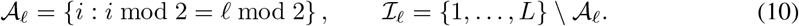

For each active residue and angular channel, the layer applies a circular rational-quadratic spline,

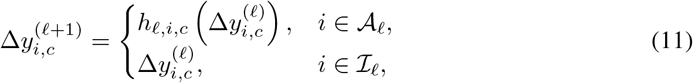

where *h*_*ℓ,i,c*_ : T → T is an invertible monotone periodic spline with *K* = 8 bins. The spline widths, heights, and derivatives are predicted by the geometry-aware conditioner.

To preserve the coupling structure, the conditioner receives the masked input

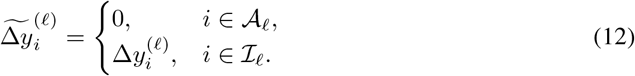

Because the variables are displacements from native torsions and ideal valence angles, zero corresponds to the reference geometry. The spline parameters for the active residues therefore depend only on the inactive variables and the conditioning inputs (*s, τ* ^nat^). The resulting Jacobian is block triangular, and inversion requires one conditioner evaluation followed by independent onedimensional spline inversions over the active variables.

To stabilize optimization, each unconstrained spline parameter *u* is soft-clipped according to

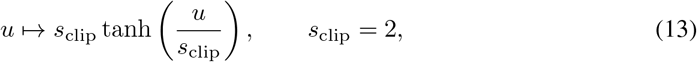

before applying the positive spline parameterization. The final spline head is zero-initialized such that every coupling layer initially represents the identity transformation.

Let

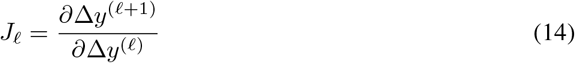

denote the Jacobian of the transformation at layer *ℓ*. Its log-determinant is

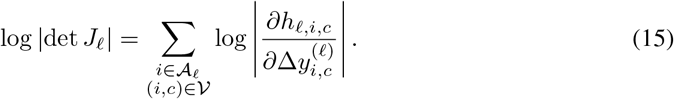

The conditional log density is therefore

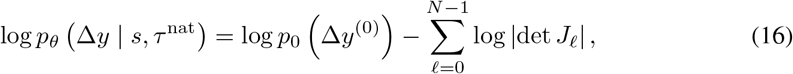

where Δ*y*^(0)^ is obtained by applying the inverse flow to Δ*y*^(*N*)^ = Δ*y*.

### 3.2 Geometry-Aware Conditioner

The conditioner predicts the circular-spline parameters for the active residues from the masked displacement input 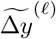 in three stages: differentiable backbone reconstruction, invariant geometric encoding, and joint sequence–structure conditioning.

#### Backbone reconstruction

The masked displacements are combined with their reference angles and reconstructed into Cartesian backbone coordinates using NeRF:

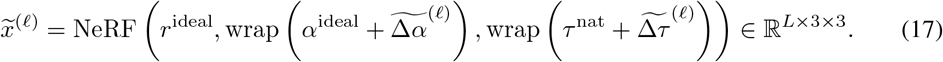

This reconstruction provides an explicit Cartesian representation of the global geometry implied by the currently observed variables.

#### Invariant geometric encoding

The reconstructed backbone is encoded by an SE(3)-invariant geometric Transformer containing *N*_geo_ = 6 frame-based, IPA-style attention blocks following the ESM3 geometric Transformer architecture Hayes et al. (2025). The encoder produces a per-residue structural representation **u**_*i*_ ∈ ℝ^*d*^. A learned binary embedding indicating whether each residue is active or inactive in the current coupling layer is then added to the encoded representation.

#### Two-track sequence–structure Transformer

The structural representations 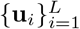 and learned amino-acid embeddings 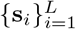 are processed by a two-track Transformer with *N*_tr_ = 6 blocks, 8 attention heads, and hidden width *d* = 256. Each block jointly attends over the sequence and structure tracks using a directional attention mask: structure tokens attend to both tracks, while sequence tokens attend only to sequence tokens. Rotary positional encodings are applied separately within each track. This design conditions the structural representation on sequence identity while preserving a geometry-independent sequence representation. Track-specific output projections and feed-forward networks are applied after attention. A linear head maps each structure token to the parameters of the six circular splines associated with that residue. For each angular channel, the head predicts *K* bin widths, *K* bin heights, and *K* periodic derivatives, yielding 18*K* spline parameters per residue:

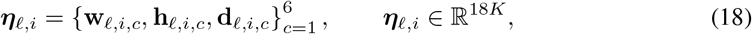

where **w**_*ℓ,i,c*_, **h**_*ℓ,i,c*_, **d**_*ℓ,i,c*_ ∈ ℝ^*K*^ denote the unconstrained bin widths, bin heights, and periodic derivatives, respectively. After soft clipping and positive parameterization, these quantities define the circular rational-quadratic spline applied to 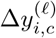:

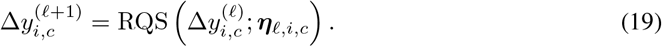

The same spline parameters determine the analytic derivative

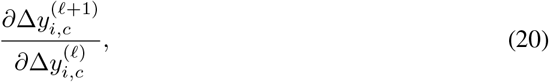

which enters the layer log-Jacobian determinant and therefore directly contributes to the conditional likelihood in Eq. 16.

### 3.3 Training Objective

A key advantage of coupling-based normalizing flows is that they support efficient one-step sampling. UniFlow exploits this property by drawing model samples during each training iteration and applying geometric supervision directly in Cartesian space.

The primary objective is the negative log-likelihood of the observed internal-coordinate displacements,

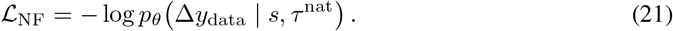

To preserve physically important structural features, we supplement this term with two sample-based geometric losses. The clash loss ℒ_clash_ penalizes nonlocal residue pairs whose generated distances fall below a prescribed threshold, while the contact loss ℒ_contact_ matches long-range contact probabilities between generated samples and the reference ensemble. The complete objective is

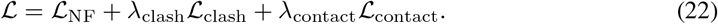

## 4 Experiments

We evaluate UniFlow for protein conformational ensemble generation across a diverse set of protein systems. Our results show that UniFlow accurately recovers equilibrium distributions from reference molecular dynamics simulations, generalizes to unseen protein systems, and enables substantially faster sampling. We further demonstrate that its explicit internal-coordinate likelihood can be converted into a differentiable energy function for stable Langevin simulations. Finally, we present comprehensive ablation studies examining the contributions of key architectural and training objective design choices.

### 4.1 Experimental Setup

#### Data

For training the sequence-conditioned normalizing flow, we use in total tens of milliseconds of equilibrium MD trajectories, which contains mdCATH Mirarchi et al. (2024) with the BioEmu simulation corpus Lewis et al. (2025), including CATH1, CATH2, Octapeptides, and MEGAsim Charron et al. (2025); Sillitoe et al. (2021); Tsuboyama et al. (2023). ATLAS Vander Meersche et al. (2024) is held out and used only as test dataset.

#### Baselines

Our primary baseline is BioEmu Lewis et al. (2025), a recent state-of-the-art model for transferable protein conformational ensemble generation trained at a scale comparable to ours. Although several diffusion- and flow-matching-based methods have been proposed for ensemble generation, BioEmu is among the few models trained on large-scale, high-quality molecular dynamics simulation data and therefore provides the most direct comparison for our setting.

We additionally compare against normalizing-flow-based baselines in the ablation study. While prior NF-based molecular ensemble generators have shown promising results, they have primarily focused on small molecules, peptides, or relatively small protein systems. To evaluate the effect of NF architecture design under our large-scale training setting, we adapt two previously proposed architectures for biomolecular modeling: PROSE Tan et al. (2026), a TARFlow-based architecture operating in Cartesian coordinates, and SplitFlow Kim et al. (2024), a related Transformer-based architecture operating in internal coordinates. These comparisons provide controlled NF baselines for assessing the contributions of UniFlow’s internal-coordinate representation, coupling architecture, and training recipe design.

### 4.2 Results

We evaluate ensemble quality by comparing generated conformational distributions against reference MD simulations. We report Jensen–Shannon divergence (JSD) between reference and generated ensembles projected onto four collective variables: end-to-end distance (EED), radius of gyration (*R*_*g*_), RMSD to the native structure, and the top two TICA components computed from *C*_*α*_ pseudo-torsion angles Schultze & Grubmuuller (2021).

We evaluate UniFlow in three settings: CATH1, a seen-protein benchmark containing 50 proteins; held-out octapeptides, which assess generalization to unseen sequences in a length-matched regime; and ATLAS, an external benchmark composed of unseen protein systems Vander Meersche et al. (2024). The results are summarized in Table 1.

**Table 1.** JS divergence against MD and wall-time on test proteins from each dataset. All timing experiments generate 2,000 i.i.d. samples on a single NVIDIA L40S GPU. UniFlow uses a batch size of 64, while BioEmu uses its default protein-length-dependent batch size.

| Quantity | CATH1 ($N = 50$) | | Octapeptide ($N = 50$) | | ATLAS ($N = 40$) | |
| --- | --- | --- | --- | --- | --- | --- |
|  | BioEmu | UniFlow | BioEmu | UniFlow | BioEmu | UniFlow |
| End-to-end | **0.062 (0.006)** | 0.093 (0.011) | 0.039 (0.000) | **0.038 (0.003)** | **0.291 (0.005)** | 0.305 (0.008) |
| $R_g$ | **0.083 (0.007)** | 0.100 (0.015) | **0.038 (0.000)** | 0.038 (0.003) | 0.323 (0.019) | **0.247 (0.012)** |
| RMSD | **0.124 (0.012)** | 0.125 (0.011) | 0.038 (0.000) | **0.037 (0.003)** | 0.366 (0.032) | **0.219 (0.017)** |
| 2D TICA | **0.182 (0.011)** | 0.190 (0.014) | 0.336 (0.005) | **0.243 (0.007)** | **0.434 (0.001)** | 0.444 (0.021) |
| Time (s) | 451.9 | **4.4** | 71.1 | **1.0** | 2181.1 | **16.2** |

UniFlow performs comparably to BioEmu across all four metrics on CATH1. On the held-out octapeptides, it achieves similar or lower JSD across the reported collective variables, demonstrating generalization to unseen peptide systems. Comparable performance is also observed on ATLAS, despite its complete exclusion from training, indicating transferability to unseen protein systems. Together, these results show that UniFlow captures both geometric observables and slow collective modes across seen and held-out protein systems.

One key advantage of UniFlow is its sampling efficiency. UniFlow generates samples by one inverse step through the normalizing flow, leading to substantially lower sampling cost. As shown in Table 1, UniFlow substantially reduces wall-clock sampling time relative to BioEmu while maintaining competitive ensemble fidelity.

### 4.3 Ablation Studies

We conduct comprehensive ablation on the normalizing flow architectures, and training objective design in Table 2. We compare four variants: TARFlow trained directly on Cartesian backbone coordinates (PROSE baseline), a Transformer-architecture circular spline flow trained on the internal coordinate (SplitFlow baseline), a UniFlow trained with only liklihood objective, and the final UniFlow model trained with both NLL and geometric objective.

**Table 2.** Ablation study of coordinate representation and coupling-mask design. We compare flow modeling directly in Cartesian space (PROSE baseline), previous flow modeling in internal coordinate (SplitFlow baseline), UniFlow trained using only the negative log-likelihood objective, and UniFlow trained using both negative log-likelihood and geometric losses.

| Dataset | Quantity | Cartesian TARFlow | SplitFlow | UniFlow (NLL only) | UniFlow |
| --- | --- | --- | --- | --- | --- |
| CATH1 | End-to-end | 0.271 (0.016) | 0.351 (0.024) | 0.263 (0.013) | **0.093 (0.011)** |
| | $R_g$ | 0.250 (0.018) | 0.415 (0.030) | 0.429 (0.027) | **0.100 (0.015)** |
|  | RMSD | 0.310 (0.023) | 0.448 (0.026) | 0.368 (0.019) | **0.125 (0.011)** |
|  | 2D TICA | 0.419 (0.012) | 0.320 (0.018) | 0.261 (0.014) | **0.190 (0.014)** |
| Octapeptide | End-to-end | 0.410 (0.010) | 0.214 (0.008) | **0.037 (0.002)** | 0.038 (0.003) |
| | $R_g$ | 0.381 (0.013) | 0.312 (0.010) | **0.035 (0.010)** | 0.038 (0.003) |
|  | RMSD | 0.480 (0.010) | 0.432 (0.013) | **0.035 (0.001)** | 0.037 (0.003) |
|  | 2D TICA | 0.595 (0.011) | 0.477 (0.031) | 0.260 (0.017) | **0.243 (0.007)** |
| ATLAS | End-to-end | 0.583 (0.023) | 0.478 (0.034) | 0.521 (0.035) | **0.305 (0.008)** |
| | $R_g$ | 0.603 (0.027) | 0.512 (0.031) | 0.497 (0.044) | **0.247 (0.012)** |
|  | RMSD | 0.627 (0.050) | 0.579 (0.049) | 0.428 (0.032) | **0.219 (0.017)** |
|  | 2D TICA | 0.680 (0.013) | 0.646 (0.023) | 0.525 (0.027) | **0.444 (0.021)** |

Cartesian TARFlow performs poorly across all datasets, indicating that direct likelihood modeling in backbone Cartesian space is difficult for coordinate-wise affine flows. SplitFlow, which instead operates in internal-coordinate space, performs similarly poorly and is particularly weak on global observables, even underperforming the Cartesian TARFlow baseline. This behavior is intuitive: likelihood optimization in internal-coordinate space is dominated by local geometric variations and does not explicitly encourage the model to capture long-range structural correlations, such as inter-strand contacts underlying *β*-sheet formation.

UniFlow addresses this limitation through a geometry-aware architecture that can propagate information across the full protein structure. However, when trained using only the negative loglikelihood objective, the optimization remains biased toward local internal-coordinate statistics. This explains why UniFlow trained with NLL alone performs comparably to, or even slightly better than, the full model on the short octapeptide systems, where local structure dominates, but degrades substantially on larger proteins whose ensembles depend more strongly on global conformational organization. The full UniFlow model achieves the best overall distributional agreement by combining the geometry-aware architecture with explicit geometric losses. These results indicate that both architectural modeling of long-range dependencies and geometric supervision are important for transferable protein ensemble generation.

### 4.4 UniFlow as a Machine-Learned Force Field for Molecular Dynamics

A key advantage of UniFlow is that it provides exact likelihood evaluation in internal-coordinate space. Let

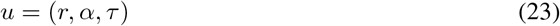

denote the complete set of backbone internal coordinates, where *r, α*, and *τ* are the bond lengths, valence angles, and torsion angles, respectively. After fixing the global translational and rotational degrees of freedom, and away from coordinate singularities, the internal coordinates provide a locally one-to-one parameterization of the corresponding Cartesian configurations:

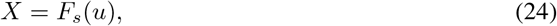

where *F*_*s*_ denotes the internal-to-Cartesian reconstruction map for a protein with sequence *s*.

Although the transferable normalizing flow is trained only on angular coordinates, its density can be extended to the complete internal-coordinate space by introducing an explicit reference distribution over bond lengths. This extension is used only for constructing the molecular-dynamics potential; direct ensemble generation continues to use ideal bond lengths.

UniFlow models the joint internal-coordinate density as

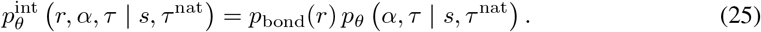

The angular factor is represented by the normalizing flow described above, with

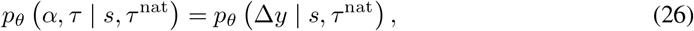

where Δ*y* contains the wrapped displacements of the valence and torsion angles from their reference values.

Because backbone bond lengths fluctuate over a substantially narrower range than the angular coordinates, we describe their density using stiff harmonic reference potentials:

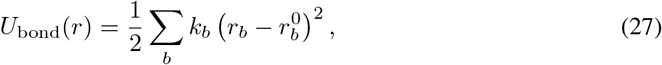

corresponding to the density with respect to the internal-coordinate measure *dr*,

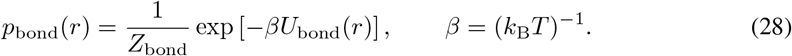

The density in Eq. equation 25 induces a properly normalized density over gauge-fixed Cartesian configurations through the geometric change of variables. Throughout this section, 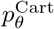 denotes a density with respect to the Euclidean volume measure on the gauge-fixed Cartesian configuration space; the six global rigid-body degrees of freedom are not included. Let

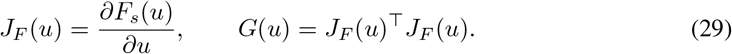

The Cartesian volume element induced by the reconstruction map is

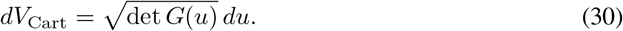

Consequently, the Cartesian density induced by UniFlow is

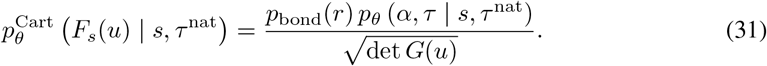

Equivalently,

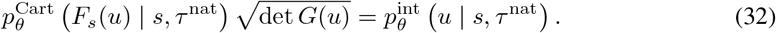

UniFlow therefore defines the explicit potential of mean force on the Cartesian coordinate

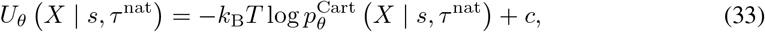

where *c* is an additive constant. Substituting Eq. equation 31 gives

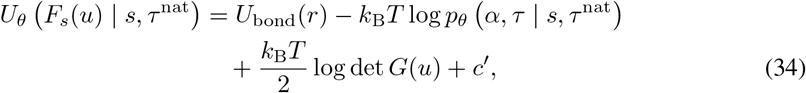

where *c*^*′*^ absorbs the normalization constant of the bond-length density. The final term accounts for the nonuniform Cartesian volume associated with the internal-coordinate parameterization. During simulation, the internal coordinates *u*(*X*) are extracted differentiably from the current Cartesian configuration. Cartesian forces are then obtained by backpropagating through the bond, angular-density, and geometric-Jacobian contributions in Eq. equation 34.

For additional stability outside regions well represented by the training distribution, we introduce a short-range repulsive potential *U*_rep_ between nonbonded atoms. The simulation potential is therefore

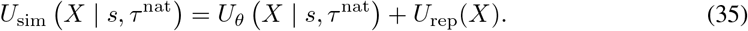

The repulsive term suppresses severe steric clashes in low-density regions and is chosen to be negligible over configurations typical of the equilibrium training distribution.

We perform Cartesian underdamped Langevin dynamics under Eq. equation 35:

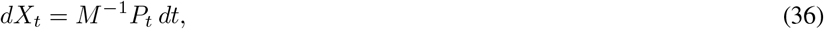

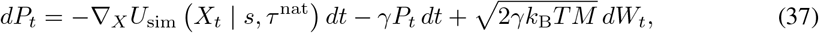

where *M* is the Cartesian mass matrix, *P*_*t*_ denotes the momenta, *γ* is the friction coefficient, and *W*_*t*_ is a standard Wiener process. The equations are integrated using the BAOAB splitting scheme.

A central challenge in developing transferable machine-learned force fields is the tension between generalization and simulation efficiency. Transferability across diverse protein systems typically requires a large and expressive model. Molecular dynamics, however, requires differentiating the potential at every integration step, making repeated force evaluation with the full transferable UniFlow model computationally expensive.

We address this limitation through system-specific distillation. Conventional machine-learned force fields generally require configurations from the target system before a distilled potential can be trained. Obtaining these configurations may itself require an external physics-based simulation or a separate generative ensemble model, reducing the practical benefit of distillation.

UniFlow avoids this requirement by serving simultaneously as a transferable ensemble generator and an explicit coarse-grained potential. Given a target protein, we first draw internal-coordinate conformations

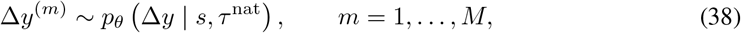

directly from the pretrained UniFlow model. These conformations are then used to train a compact system-specific student flow using the same negative-log-likelihood and geometric objectives as the transferable model. Because the student only needs to represent one protein rather than generalize across sequences and structures, the sequence-conditioning pathway and geometry-aware Transformer are no longer required. We therefore replace the conditioner in each coupling layer with a lightweight multilayer perceptron, yielding a substantially smaller model for repeated force evaluation during molecular dynamics.

We visualize the free-energy landscapes obtained from UniFlow-driven Langevin dynamics in Fig. 2. The two-dimensional TICA free-energy surfaces show that UniFlow MD recovers the major metastable basins observed in the reference trajectories. The one-dimensional free-energy profiles along end-to-end distance, radius of gyration, and RMSD further show that the simulations capture the dominant low-free-energy regions well.

**Figure 2.**
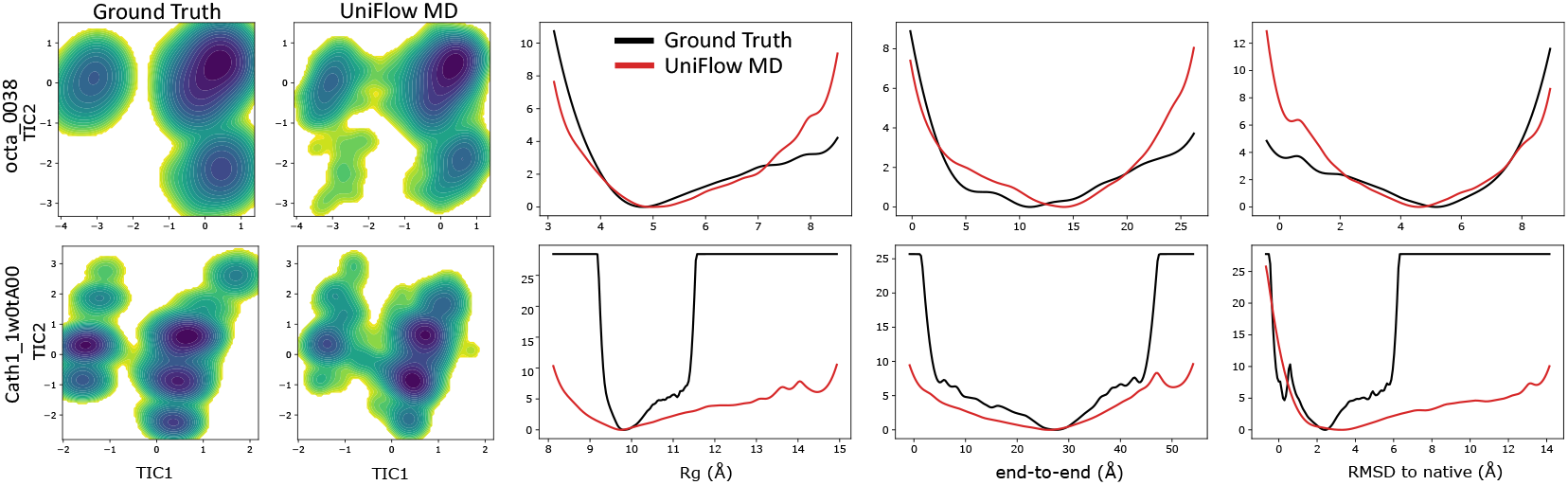
Free-energy landscapes obtained from UniFlow-driven Langevin dynamics. The top row shows an octapeptide system, and the bottom row shows a CATH1 protein. For each system, 64 parallel walkers were simulated for 50 ns each, corresponding to an aggregate simulation time of 3.2 *µ*s per system.

Simulation efficiency is reported in Table 3. Direct simulation with the pretrained UniFlow model is expensive because every force evaluation requires the geometry-aware Transformer and sequential NeRF conditioner. Distillation into a system-specific MLP substantially reduces this cost. Combining distillation with `torch.compile` decreases the wall-clock time of a 2 ns simulation with 64 parallel walkers from 134.2 to 5.3 hours for a 61-residue CATH1 protein and from 128.77 to 5.2 hours for an eight-residue peptide.

**Table 3.** Wall-clock time in hours for a 2 ns Langevin simulation (10^6^ integration steps) with 64 parallel walkers and a timestep of *dt* = 2 fs. Distillation replaces the geometry-aware Transformer and sequential NeRF conditioner with a lightweight system-specific MLP. Combining distillation with `torch.compile` provides the largest speedup. Lower is better.

| Method | CATH1-0 (61 residues) | Octapeptide-0 (8 residues) |
| --- | --- | --- |
| Pretrained UniFlow | 134.2 | 128.77 |
| Pretrained UniFlow + `torch.compile` | 66.4 | 36.9 |
| Distilled UniFlow | 45.7 | 44.6 |
| Distilled UniFlow + `torch.compile` | **5.3** | **5.2** |

## 5 Discussion

### Limitations

UniFlow is reference-conditioned and requires a reference backbone torsion configuration at generation time. The reference provides a structural frame for the angular displacement representation, but does not restrict the model to small local fluctuations around a single conformation. UniFlow can still represent broad and multimodal conformational distributions when such states are supported by the training data. Nevertheless, the generated distribution may depend on the supplied reference. Extending UniFlow to reference-free conditioning could reduce this dependence and improve coverage of highly heterogeneous conformational landscapes.

### Opportunities

The combination of exact likelihood evaluation, differentiable energy computation, and efficient sampling creates several opportunities beyond ensemble generation. In particular, UniFlow can be combined with physics-based energy functions through variational force matching, providing a route toward hybrid coarse-grained force fields that integrate learned equilibrium distributions with explicit physical interactions.

UniFlow’s rapid i.i.d. generation also makes it practical to fine-tune the learned distribution against large-scale datasets of global molecular properties. Quantities such as folding stability, mutation-induced free-energy changes, structural compactness, and other experimental observables can be evaluated over generated conformations and incorporated through differentiable objectives, following the same principle demonstrated by the geometric loss.

## References

Marloes Arts, Victor Garcia Satorras, Chin-Wei Huang, Daniel Zugner, Marco Federici, Cecilia Clementi, Frank Noé, Robert Pinsler, and Rianne van den Berg. Two for one: Diffusion models and force fields for coarse-grained molecular dynamics. Journal of Chemical Theory and Computation, 19(18):6151–6159, 2023.

Nicholas E Charron, Klara Bonneau, Aldo S Pasos-Trejo, Andrea Guljas, Yaoyi Chen, Félix Musil, Jacopo Venturin, Daria Gusew, Iryna Zaporozhets, Andreas Krämer, et al. Navigating protein landscapes with a machine-learned transferable coarse-grained model. Nature chemistry, pp. 1–9, 2025.

Laurent Dinh, Jascha Sohl-Dickstein, and Samy Bengio. Density estimation using real nvp. arXiv preprint arXiv:1605.08803, 2016.

Yuanqi Du, Michael Plainer, Rob Brekelmans, Chenru Duan, Frank Noe, Carla P Gomes, Alan Aspuru-Guzik, and Kirill Neklyudov. Doob’s lagrangian: A sample-efficient variational approach to transition path sampling. Advances in Neural Information Processing Systems, 37:65791–65822, 2024.

Conor Durkan, Artur Bekasov, Iain Murray, and George Papamakarios. Neural spline flows. Advances in neural information processing systems, 32, 2019.

Aleksander EP Durumeric, Yaoyi Chen, Aldo S Pasos-Trejo, Frank Noé, and Cecilia Clementi. Learning data-efficient coarse-grained molecular dynamics from forces and noise. Nature Communications, 2026.

Xiang Fu, Tian Xie, Nathan J Rebello, Bradley D Olsen, and Tommi Jaakkola. Simulate time-integrated coarse-grained molecular dynamics with multi-scale graph networks. arXiv preprint arXiv:2204.10348, 2022.

Thomas Hayes, Roshan Rao, Halil Akin, Nicholas J Sofroniew, Deniz Oktay, Zeming Lin, Robert Verkuil, Vincent Q Tran, Jonathan Deaton, Marius Wiggert, et al. Simulating 500 million years of evolution with a language model. Science, 387(6736):850–858, 2025.

Bowen Jing, Bonnie Berger, and Tommi Jaakkola. Alphafold meets flow matching for generating protein ensembles. arXiv preprint arXiv:2402.04845, 2024a.

Bowen Jing, Hannes Stärk, Tommi Jaakkola, and Bonnie Berger. Generative modeling of molecular dynamics trajectories. Advances in Neural Information Processing Systems, 37:40534–40564, 2024b.

John Jumper, Richard Evans, Alexander Pritzel, Tim Green, Michael Figurnov, Olaf Ronneberger, Kathryn Tunyasuvunakool, Russ Bates, Augustin Žídek, Anna Potapenko, et al. Highly accurate protein structure prediction with alphafold. nature, 596(7873):583–589, 2021.

Joseph C Kim, David Bloore, Karan Kapoor, Jun Feng, Ming-Hong Hao, and Mengdi Wang. Scalable normalizing flows enable boltzmann generators for macromolecules. arXiv preprint arXiv:2401.04246, 2024.

Tony Lelièvre, Geneviève Robin, Innas Sekkat, Gabriel Stoltz, and Gabriel Victorino Cardoso. Generative methods for sampling transition paths in molecular dynamics. ESAIM: Proceedings and Surveys, 73:238–256, 2023.

Sarah Lewis, Tim Hempel, José Jiménez-Luna, Michael Gastegger, Yu Xie, Andrew YK Foong, Victor García Satorras, Osama Abdin, Bastiaan S Veeling, Iryna Zaporozhets, et al. Scalable emulation of protein equilibrium ensembles with generative deep learning. Science, pp. eadv9817, 2025.

Yikai Liu, Haoyang Zheng, Lining Mao, Yanbin Wang, Ming Chen, and Guang Lin. Protdyn: a foundation protein language model for thermodynamics and dynamics generation. arXiv preprint arXiv:2510.00013, 2025.

Yikai Liu, Guang Lin, and Ming Chen. Conformflow: scalable normalizing flow for protein conformational ensemble generation. bioRxiv, pp. 2026–06, 2026.

Jiarui Lu, Bozitao Zhong, Zuobai Zhang, and Jian Tang. Str2str: A score-based framework for zero-shot protein conformation sampling. arXiv preprint arXiv:2306.03117, 2023.

Maciej Majewski, Adrià Pérez, Philipp Thölke, Stefan Doerr, Nicholas E Charron, Toni Giorgino, Brooke E Husic, Cecilia Clementi, Frank Noé, and Gianni De Fabritiis. Machine learning coarsegrained potentials of protein thermodynamics. Nature communications, 14(1):5739, 2023.

Antonio Mirarchi, Toni Giorgino, and Gianni De Fabritiis. mdcath: A large-scale md dataset for data-driven computational biophysics. Scientific Data, 11(1):1299, 2024.

Frank Noé, Simon Olsson, Jonas Köhler, and Hao Wu. Boltzmann generators: Sampling equilibrium states of many-body systems with deep learning. Science, 365(6457):eaaw1147, 2019.

George Papamakarios, Theo Pavlakou, and Iain Murray. Masked autoregressive flow for density estimation. Advances in neural information processing systems, 30, 2017.

Sanjeev Raja, Martin Šípka, Michael Psenka, Tobias Kreiman, Michal Pavelka, and Aditi S Krishnapriyan. Action-minimization meets generative modeling: Efficient transition path sampling with the onsager-machlup functional. arXiv preprint arXiv:2504.18506, 2025.

Paul Robustelli, Stefano Piana, and David E Shaw. Developing a molecular dynamics force field for both folded and disordered protein states. Proceedings of the National Academy of Sciences, 115 (21):E4758–E4766, 2018.

Steffen Schultze and Helmut Grubmuuller. Time-lagged independent component analysis of random walks and protein dynamics. Journal of Chemical Theory and Computation, 17(9):5766–5776, 2021.

David E Shaw, Paul Maragakis, Kresten Lindorff-Larsen, Stefano Piana, Ron O Dror, Michael P Eastwood, Joseph A Bank, John M Jumper, John K Salmon, Yibing Shan, et al. Atomic-level characterization of the structural dynamics of proteins. Science, 330(6002):341–346, 2010.

Ian Sillitoe, Nicola Bordin, Natalie Dawson, Vaishali P Waman, Paul Ashford, Harry M Scholes, Camilla SM Pang, Laurel Woodridge, Clemens Rauer, Neeladri Sen, et al. Cath: increased structural coverage of functional space. Nucleic acids research, 49(D1):D266–D273, 2021.

Charlie Tan, Majdi Hassan, Leon Klein, Saifuddin Syed, Dominique Beaini, Michael Bronstein, Alexander Tong, and Kirill Neklyudov. Amortized sampling with transferable normalizing flows. Advances in Neural Information Processing Systems, 38:94290–94325, 2026.

Kotaro Tsuboyama, Justas Dauparas, Jonathan Chen, Elodie Laine, Yasser Mohseni Behbahani, Jonathan J Weinstein, Niall M Mangan, Sergey Ovchinnikov, and Gabriel J Rocklin. Mega-scale experimental analysis of protein folding stability in biology and design. Nature, 620(7973):434–444, 2023.

Yann Vander Meersche, Gabriel Cretin, Aria Gheeraert, Jean-Christophe Gelly, and Tatiana Galochkina. Atlas: protein flexibility description from atomistic molecular dynamics simulations. Nucleic acids research, 52(D1):D384–D392, 2024.

Hannah K Wayment-Steele, Adedolapo Ojoawo, Renee Otten, Julia M Apitz, Warintra Pitsawong, Marc Hömberger, Sergey Ovchinnikov, Lucy Colwell, and Dorothee Kern. Predicting multiple conformations via sequence clustering and alphafold2. Nature, 625(7996):832–839, 2024.

James C Whisstock and Arthur M Lesk. Prediction of protein function from protein sequence and structure. Quarterly reviews of biophysics, 36(3):307–340, 2003.

Shuangfei Zhai, Ruixiang Zhang, Preetum Nakkiran, David Berthelot, Jiatao Gu, Huangjie Zheng, Tianrong Chen, Miguel Angel Bautista, Navdeep Jaitly, and Josh Susskind. Normalizing flows are capable generative models. arXiv preprint arXiv:2412.06329, 2024.

Shuxin Zheng, Jiyan He, Chang Liu, Yu Shi, Ziheng Lu, Weitao Feng, Fusong Ju, Jiaxi Wang, Jianwei Zhu, Yaosen Min, et al. Predicting equilibrium distributions for molecular systems with deep learning. Nature Machine Intelligence, 6(5):558–567, 2024.

